# *In vivo* Mapping of Cellular Resolution Neuropathology in Brain Ischemia by Diffusion MRI

**DOI:** 10.1101/2023.08.08.552374

**Authors:** Dan Wu, Victoria Turnbill, Hong-Hsi Lee, Xiaoli Wang, Ruicheng Ba, Piotr Walczak, Lee J. Martin, Els Fieremans, Dmitry S. Novikov, Frances J. Northington, Jiangyang Zhang

## Abstract

Non-invasive mapping of cellular pathology can provide critical diagnostic and prognostic information. Recent developments in diffusion MRI have produced new tools for examining tissue microstructure at a level well below the imaging resolution. Here, we report the use of diffusion time (*t*)-dependent diffusion kurtosis imaging (*t*DKI) to simultaneously assess the morphology and transmembrane permeability of cells and their processes in the context of pathological changes in hypoxic-ischemic brain (HI) injury. Through Monte Carlo simulations and cell culture organoid imaging, we demonstrate feasibility in measuring effective size and permeability changes based on the peak and tail of *t*DKI curves. In a mouse model of HI, *in vivo* imaging at 11.7T detects a marked shift of the *t*DKI peak to longer *t* in brain edema, suggesting swelling and beading associated with the astrocytic processes and neuronal neurites. Furthermore, we observed a faster decrease of the *t*DKI tail in injured brain regions, reflecting increased membrane permeability that was associated with upregulated water exchange upon astrocyte activation at acute stage as well as necrosis with disrupted membrane integrity at subacute stage. Such information, unavailable with conventional diffusion MRI at a single *t,* can predict salvageable tissues. For a proof-of-concept, *t*DKI at 3T on an ischemic stroke patient suggested increased membrane permeability in the stroke region. This work therefore demonstrates the potential of *t*DKI for *in vivo* detection of the pathological changes in microstructural morphology and transmembrane permeability after ischemic injury using a clinically translatable protocol.

## INTRODUCTION

Disrupted cellular structure and membrane integrity are common indicators of acute tissue pathophysiology and injury in the brain. For instance, swelling of cell bodies and processes and disruptions of membrane structures are hallmarks of ischemic stroke. Currently, assessment of neuropathology at the cellular level mostly relies on histopathology, which is mostly postmortem and limited both spatially and temporally. *In vivo* non-invasive mapping of the brain pathology in real time, if available, will lead to accurate diagnosis and prognosis (e.g., tissue survival) to enable more effective and personalized treatments.

Magnetic resonance imaging (MRI) provides non-invasive tools to assess brain structure and physiology. Particularly, the development of diffusion MRI (dMRI) utilizes diffusion of water molecules to probe tissue microstructure ^1–4^, and the acquired dMRI signals reflect ensemble-averaged *μ*m-level tissue microstructural features well-below the spatial resolution of MRI (mm-level). A prime example is the use of dMRI to detect acute ischemic stroke^5^, where the dramatic drop in the water diffusivities within minutes after ischemic insult ^6^ has been linked to tissue microstructural changes^7–10^, even though the exact pathology (e.g., neurons or astrocytes, cell bodies or processes) that contribute to the drop are unresolved. The diagnostic power of dMRI may be enhanced if it can be linked to specific cellular pathology.

A series of work in diffusion time (*t*)-dependent dMRI ^10–13^ offers unprecedented opportunity to assess evolving cellular pathology. The measured *t-*dependency is used to infer the spatial scale and statistical spatial distributions of tissue microstructure barriers to water diffusion ^14–18^. This technique has proven useful for interrogating ischemic brain injuries as reported by us and others^8,10,19–21^. *t*-dependency of dMRI signals also reflects transmembrane water exchange^13,22–29^, regulated by active and passive water channels located on cell membranes, and thus, can be linked to metabolism and membrane integrity^22,29^. If the microstructural and exchange effects have distinct time scales^28,30^, they can be in principle measured separately by *t*-dependent dMRI^25,26^. In the case of ischemic brain injury, this may provide critical insights into the dynamics of cellular swelling, beading, and membrane disruption^7–10^. For instance, astrocytic activation induces increased water fluxes via up-regulated aquaporin-4 (AQP4) channels ^31^ and also restructures the astrocytic processes to tune the volume ^32^, and thus, inflicting both microstructural size and membrane permeability change.

In this study, we used *t*-dependent diffusion kurtosis imaging (*t*DKI), which measures the *t-* dependent changes of the non-Gaussian diffusion component in dMRI signals, to simultaneously infer microstructural size and membrane integrity via water exchange (**Fig. 1**). We hypothesized that these new markers are sensitive to cellular-level pathological changes after ischemic brain injury and can help identify subacutely salvageable tissue. We carried out numerical simulations, *in vitro* experiments, and *in vivo* experiments in a mouse model of neonatal hypoxia-ischemia (HI) to test the hypothesis, followed by a proof-of-concept study in an ischemic stroke patient.

**Fig. 1:**
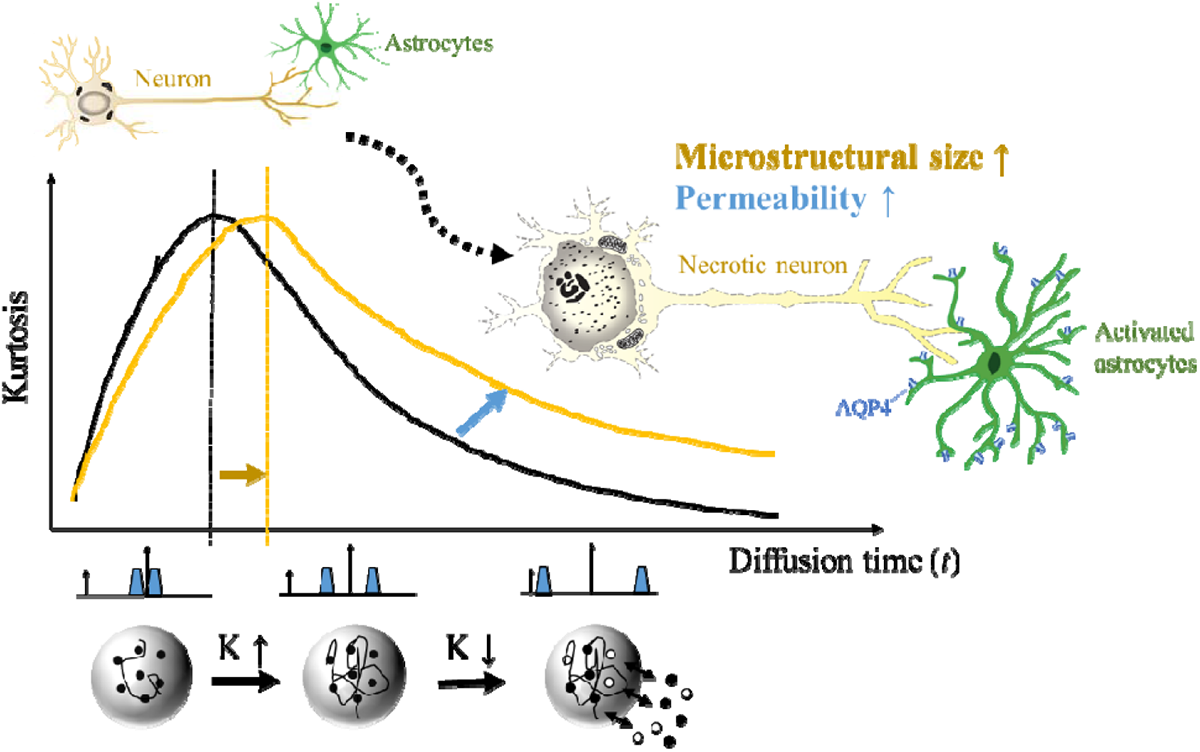
Schematics of *t*DKI in measuring microstructural size and membrane integrity. Normal neuron and astrocyte morphology shown in upper left cartoon. Diffusion kurtosis (K) in biological tissues changes non-monotonically as *t* increases, with an initial rise attributed to the water-restrictive effects of tissue barriers, followed by a descent as water molecules exchange between intra- and extra-cellular spaces. We hypothesize that the peak *t* (*t*_peak_)of the *t*DKI curve indicates the ensemble-averaged size of tissue barriers, or effective structure size, while the post-peak descending curve reflects membrane permeability due to water redistribution, thereby enabling the assessment of key neuropathological features following hypoxic-ischemic stroke-like brain injuries, including neuronal swelling and membrane disruption, astrocytic activation and fragmentary degeneration, and increase of transmembrane water exchange due to AQP4-channeled flux.

## RESULTS

### Simulations to investigate the effects of microstructural size and permeability on tDKI for cell body and processes

Monte Carlo simulations ^33,34^ based on randomly packed cylinders (**Fig. 2a,b**) and spheres (**Fig. 2c,d**), representing simplified cell processes (neurites/astrocytic processes, diameters of 1.0-1.4 µm) and cell bodies (diameters 10-14 µm), respectively, demonstrated the behaviors of *t*-dependent diffusivity and kurtosis in response to changes in structural size and membrane permeability. The monotonically decreased diffusivities with increasing *t* (**Fig. 2a,c**) is the result of restrictive effects from microstructures including the so-called coarse-graining process (i.e. for water molecules to experience the restrictive effects of structural barriers across multiple scales) ^3^. Diffusion kurtosis, in contrast, demonstrated a non-monotonical change (**Fig. 2b,d**) with *t*. Particularly, the peak *t* (*t*_peak_) was < 5 milliseconds (ms) for the cell processes phantom (**Fig. 2b**) and 20-35 ms for the cell body phantom (**Fig. 2d**), with increasing structural size shifting the peaks towards longer *t*.

The effect of permeability on *t*_peak_ depended on the relative time scales of coarse-graining and water exchange. For the cell processes phantom, the time for completing the coarse graining (<5 ms) was shorter than the transmembrane exchange times (*τ_ex_*) used in the simulations (10-200 ms), and the *t*_peak_ mostly depended on diameter rather than *τ_ex_* (**Suppl. Table S1**). For the cell body phantom, the range of *τ_ex_* overlapped with the time for completing the coarse graining (∼50 ms), and thereby, the *t*_peak_ was modulated by both cell size and permeability (**Suppl. Table S1**).

*τ_ex_*, on the other hand, had a large effect on the asymptotic decrease of kurtosis with *t*, with faster decrease associated with lower *τ_ex_*. By fitting the Kärger model^26,35^ to the simulated *t*DKI data, the estimate *τ_ex_* matched ground truth for the cell processes phantom (**Suppl. Table S2**) but showed large deviations for the cell phantom, due to residual restrictive effects from microstructures confounded the estimation of *τ_ex_*.

Under pathological conditions, the morphology of cell bodies and processes as well as membrane permeability may change simultaneously. The simulation results pointed out that decoupling the effects of size and permeability from *t*DKI, without considering structural disorder ^10^, is possible for cell processes with small microstructural scale (1-2 µm), with the *t*_peak_ related to the size of the restrictions and decreasing tail of *t*DKI curve reflecting the transmembrane water exchange, respectively.

**Fig. 2:**
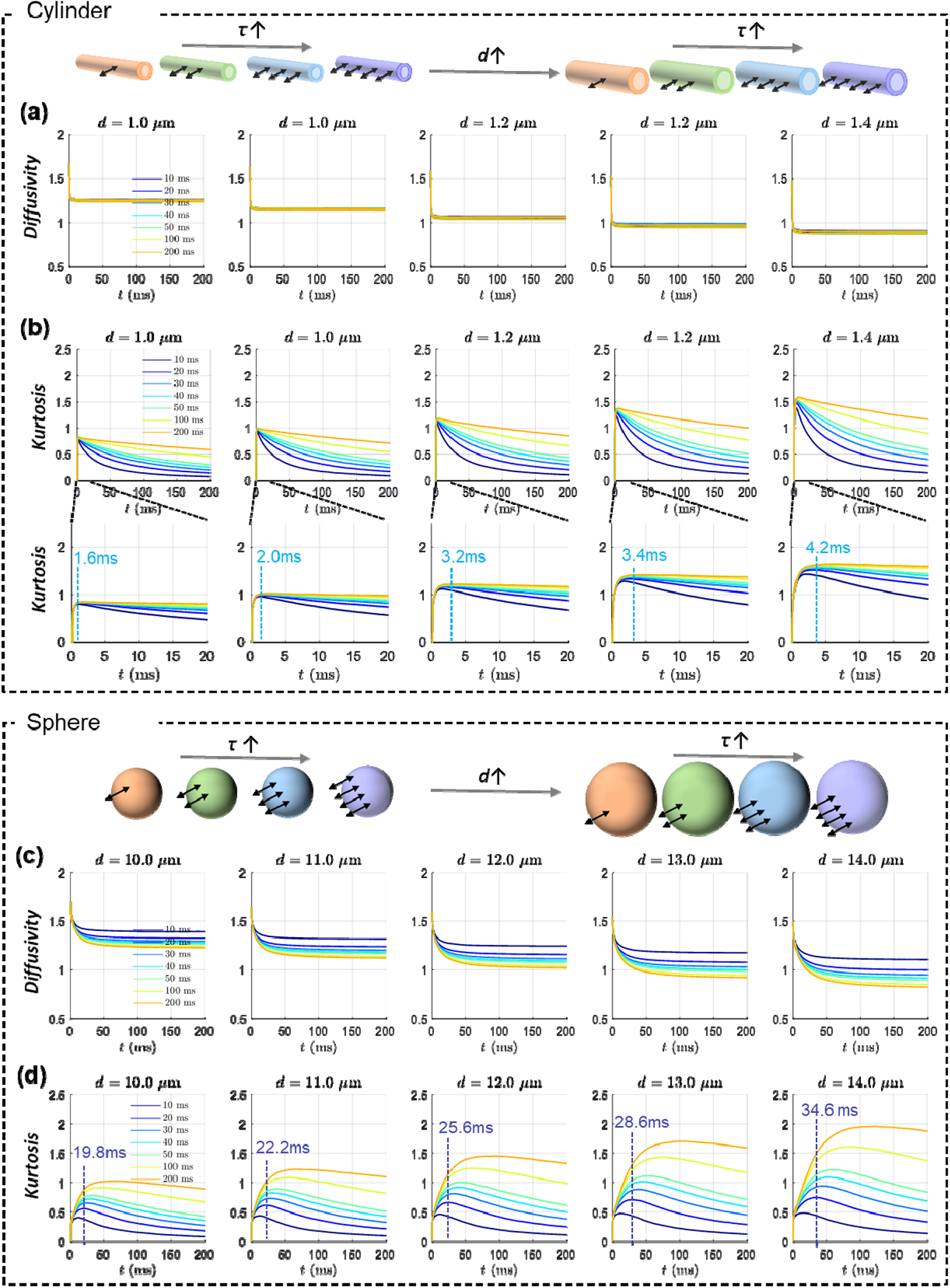
Monte-Carlo simulations of the *t*-dependent changes of the diffusivity and kurtosis at varying structure sizes and transmembrane exchange times (*τ*_ex_). (**a-b**) Simulated diffusivity (top row) and kurtosis (bottom two rows) at *t* range of 0-200 milliseconds (ms) for the cylinder phantom that represents cell processes with diameters ranging from 1.0 to 1.4 µm and τ_ex_ of 10-200 ms. The intracellular volume fraction was set to 0.3 for the 1 µm case and increased with the diameter. While the diffusivity dropped rapidly and stabilized for *t* beyond 10 ms, the kurtosis first increased and subsequently decreased (see zoom-in views, second row in **b**, for the rapid rise of kurtosis) with *t*_peak_ shifting from 1.6 to 4.2 ms (at *τ*_ex_ =50ms, typical for rodent gray matter^28^) as the diameter increased. The kurtosis further decreased with *t* and shorter τ_ex_ led to fast decay of the kurtosis. Bidirectional arrows in the cartoon the indicate permeability. (**c-d**) Simulated diffusivity (top row) and kurtosis (bottom row) for the sphere phantom that represents cell bodies with diameters ranging from 10 to 14 µm and τ_ex_ of 10-200 ms. The intracellular volume fraction was set to 0.3 for the 10 µm case and increased with the diameter. The diffusivity gradually decreased and stabilized for *t* beyond 50 ms, and kurtosis showed a non-monotonic change with peak positions increasing from 19.8 to 34.6 ms (at *τ*_ex_ =20ms, typical for gray matter^28^) as the diameter increased. Similarly, shortening the *τ*_ex_ resulted in fast decay of the kurtosis tail.

### In vitro tDKI in cultured brain organoids showed the feasibility of mapping structural size and membrane permeability

In real brain tissues with a mixture of neurons, glial cells, and their processes, can we extract information on effective structural size and membrane permeability from *t*DKI? To answer this question, we imaged 3D forebrain-specific organoids differentiated from human induced pluripotent stem cells ^36^, which resemble the architecture composition and physiology of forebrain. *In vitro* scans of the fixed organoids were performed on an 11.7 Tesla NMR spectrometer using both pulsed gradient spin-echo (PGSE) and stimulated echo acquisition mode (STEAM) dMRI sequences (**Suppl. Fig. S1**) to cover a wide range of *t* from 7-200 ms. The organoid core contained dead tissue/cells with membranes lacking integrity and digested, and the organoid rim contained more living tissue/cells with more intact membranes before fixation, as illustrated in Fig. 3g. To simulate the effects of changing structure sizes, we acquired *t*DKI data with temperatures gradually increased from 300K to 320K (300K≈ room temperature, 310K≈ body temperature). Because water diffusivity (*D*) increases with temperature, increasing temperature will increase the diffusion length scale 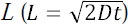, or equivalently, scale down the relative microstructure size experienced by the water molecules (**Fig. 3a**). As the temperature increased from 300K to 320K, the diffusivity increased overall as expected, and the monotonic decrease with *t* was preserved (**Fig. 3b**).

The *t*DKI curve measured at 300K in the core area had a *t*_peak_ around 20 ms (**Fig. 3c**). As the temperature rose, the kurtosis peak gradually shifted to shorter *t* (e.g. 15 ms at 305K) and eventually below 7 ms (the shortest *t* used) at temperatures above 310K (body temperature equivalent). Similar *t*-dependent changes were also observed in the rim region. This temperature-dependent behavior of kurtosis agreed with the simulation results in **Fig. 2** as the decreased effective structure size should shift the kurtosis peak towards shorter *t*.

The *t*_peak_ of the *t*DKI curve observed in the organoid provided clues on the effective structure size in the organoids. The *t*_peak_ ≤10 ms at close to body temperature (**Fig. 3e**) suggested an effective structure size that was closer to the size of cell processes (1-2 µm) than those of cell bodies. This was not surprising as light microscopy data have shown that the relative volume of neurites (dendrites & axons) and glial processes were about 4 times the volumes of somas^37^. The maps of *t*_peak_ showed that the core had higher *t*_peak_ than those of the rim (**Fig. 3d,e**), indicating larger effective structure sizes there, which may be explained by less densely populated neurites and glial processes in the rim than the core shown by the SMI-31 and GFAP immunocytochemistry (**Fig. 3g**).

As the diffusivities remained largely unchanged for *t* > 20 ms, suggesting completed coarse graining, the observed *t*DKI tails mainly reflected water exchange. It was also observed that the descending tail of the kurtosis became steeper at higher temperatures. Fitting the Kärger model to the *t*DKI data, the estimated *τ*_ex_ was considerably reduced as the temperature increased (*n*=8 samples, **Fig. 3f**), likely due to increased water mobility and/or higher membrane permeability at higher temperature. In addition, significantly lower *τ*_ex_ (higher permeability) was observed in the core than the rim (**Fig. 3f**), which agreed with the necrotic cell pathology in the core as shown in the H&E and STEM121 staining (**Fig. 3g**), suggesting disrupted membrane integrity during necrosis. The organoid data suggested that the effective structure size in brain tissue was closer to the size of cellular processes (e.g. glial processes and axons) and separation of structure size and exchange effects from *t*DKI signals was indeed possible.

**Fig. 3:**
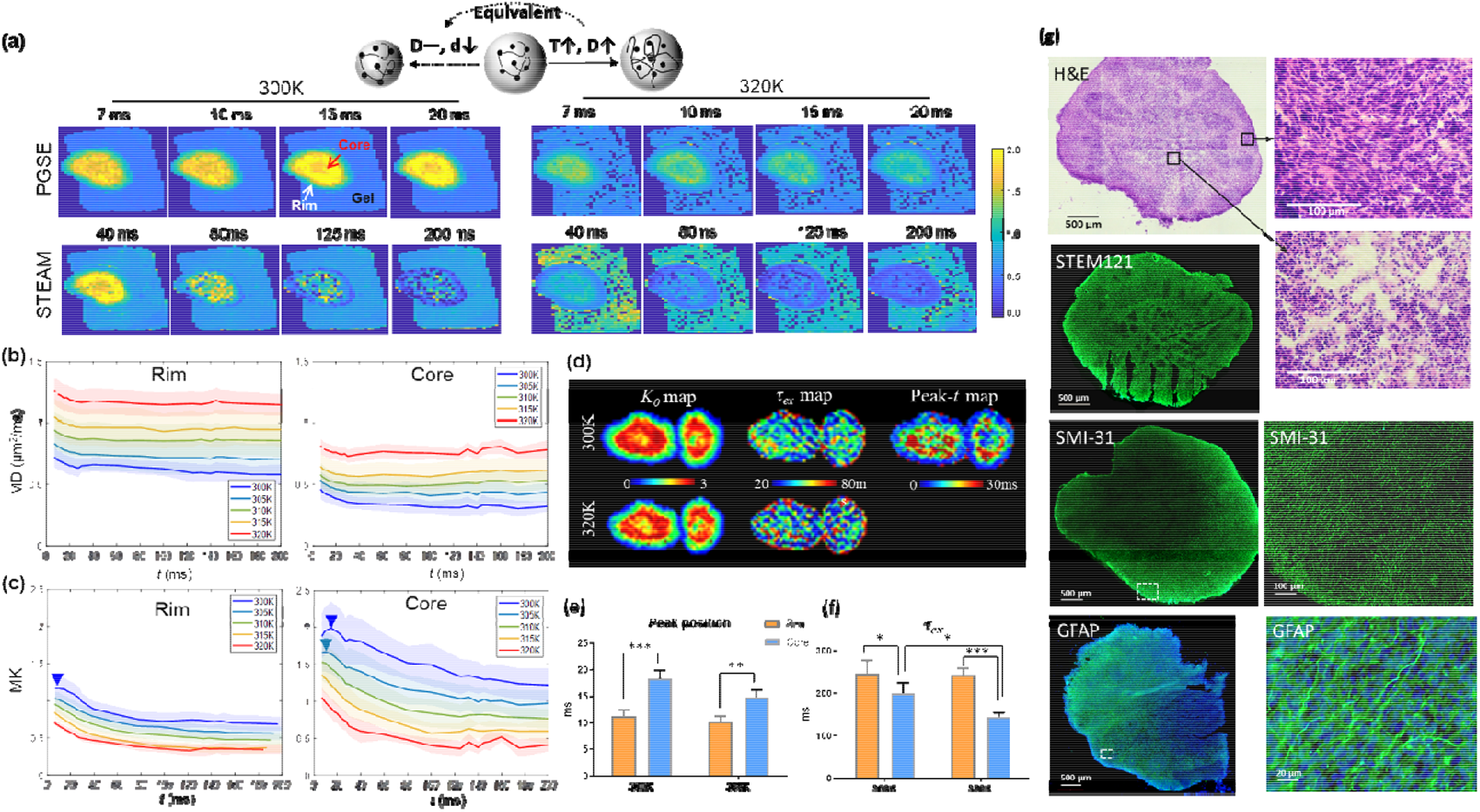
*t*DKI of the *in vitro* organoids. (**a**) Mean kurtosis maps of an organoid at varying *t*s from 7-200 ms at two temperature settings of 300K and 320K, using PGSE (for 7-20ms) and STEAM (for 40-200ms) dMRI sequences. Here, increasing temperature (*T*) has the equivalent effect as reducing microstructural size on dMRI signals. (**b-c**) *t*-dependent mean diffusivity (MD) and mean kurtosis (MK) curves at five different temperatures, obtained in the core and rim regions from eight organoid samples with the shadows showing the standard deviations. The MK curves in the core region showed a peak around 20 ms (blue arrow) at 300K, which shifted to shorter *t* as temperature increased. (**d**) Voxel-wise *K*_0_ map and τ_ex_ map as fitted from the Kärger model and *t*_peak_ maps of the organoids scanned at temperatures of 300K and 320K. (**e**) Statistical comparison of the *t*_peak_ detected in the core and rim regions at 300K and 305K (*n*=8). The peaks shifted below 7 ms above 305K. (**f**) Statistical comparison of the *t*_peak_ detected in the core and rim regions at 300K and 320K. \**p*<0.05, \*\**p*<0.01, \*\*\**p*<0.001 by paired t-test. (**g**) H&E, STEM121, SMI-31, and glial fibrillary acidic protein (GFAP) staining of the organoid. The zoomed-in views of H&E indicated necrotic changes with pyknotic nuclei and eosinophilic cell bodies in the core region, and the increased SMI-31and GFAP expression in the rim region indicated the presence of axons and astrocytes, greater than that seen in the core.

### tDKI highlights morphological changes in the cell processes upon HI injury

Taking into account the data from simulations and *in vitro* experiments of organoids support the concept of *t*DKI can detect the size and membrane integrity of cellular processes. We next tested tDKI on a widely used mouse model of neonatal HI injury (unilateral ligation of the carotid artery followed by hypoxia). This model exhibits a spectrum of ipsilateral stroke-like pathophysiology, including neuronal swelling, neurite beading, astrocytic activation, and necrotic and apoptotic cell death and the contralateral hemisphere can serve a non-stroke comparator^38,39^. In about one third of the mice (13 out of 41, see **Suppl. Table S3**), severe edema in the ipsilateral cortex and hippocampus was observed, as early as 3hrs post HI as indicated by decreased diffusivity and increased kurtosis (approximately 25% and 100% with respect to the contralateral side, respectively) and hyperintense T_2_-signals (**Fig. 4a,b**).

In the edema region, diffusivity showed higher *t*-dependency compared to the contralateral side (**Suppl. Fig. S2a-d**), similar to our previously reports based on this model^19,20^, and *t*DKI curve measured using PGSE for *t* in the 7-40 ms range only showed the descending portion (**Suppl. Fig. S2e-h**), suggesting that *t*_peak_ was shorter than 7ms. To capture the *t*DKI peak, we used the bipolar pulsed gradient (BPG) diffusion encoding (**Suppl. Fig. S1**) to acquire dMRI signals with *t* of 3-10 ms, and the observed peak appeared around 7-8 ms in the edema regions (e.g. the hippocampal CA1 region in **Fig. 4c**). The *t*DKI peak was not evident on the contralateral side, suggesting that normal tissue had a *t*_peak_ below 3 ms. Voxel-wise mapping of the *t*_peak_ in a severely injured mice further illustrated increased *t*_peak_ in the ipsilateral cortex and hippocampus at 3hrs after injury (**Fig. 4d**). The range of *t*_peak_ (**Fig. 4e**) again suggested that effective structural size in the mouse brain was comparable to cell processes, and the shift of peak position to longer *t* in edema may reflect volume expansion, including beading, of the astrocytic processes and neurites upon HI. This interpretation is supported by the GFAP immunohistochemical staining at 3hrs after HI (**Fig. 4f**). The ipsilateral hippocampus shows enrichment of GFAP immunoreactivity overall compared to the contralateral hippocampus., which is particularly striking in the CA1 region. The enrichment appears to be due to intensified GFAP immunoreactivity in ipsilateral individual swollen astrocytes and beading of their processes in the neuropil of the stratum oriens and stratum pyramidale (**Fig. 4f I’, II’**). Moreover, neurofilament staining indicates axonal beading in the stratum pyramidale and stratum radiata of the hippocampus (**Fig. 4g**) at 3hrs. The *t*DKI peak was less evident at 48hrs (**Fig. 4c**), indicating the beading phenomenon may be transient.

**Fig. 4:**
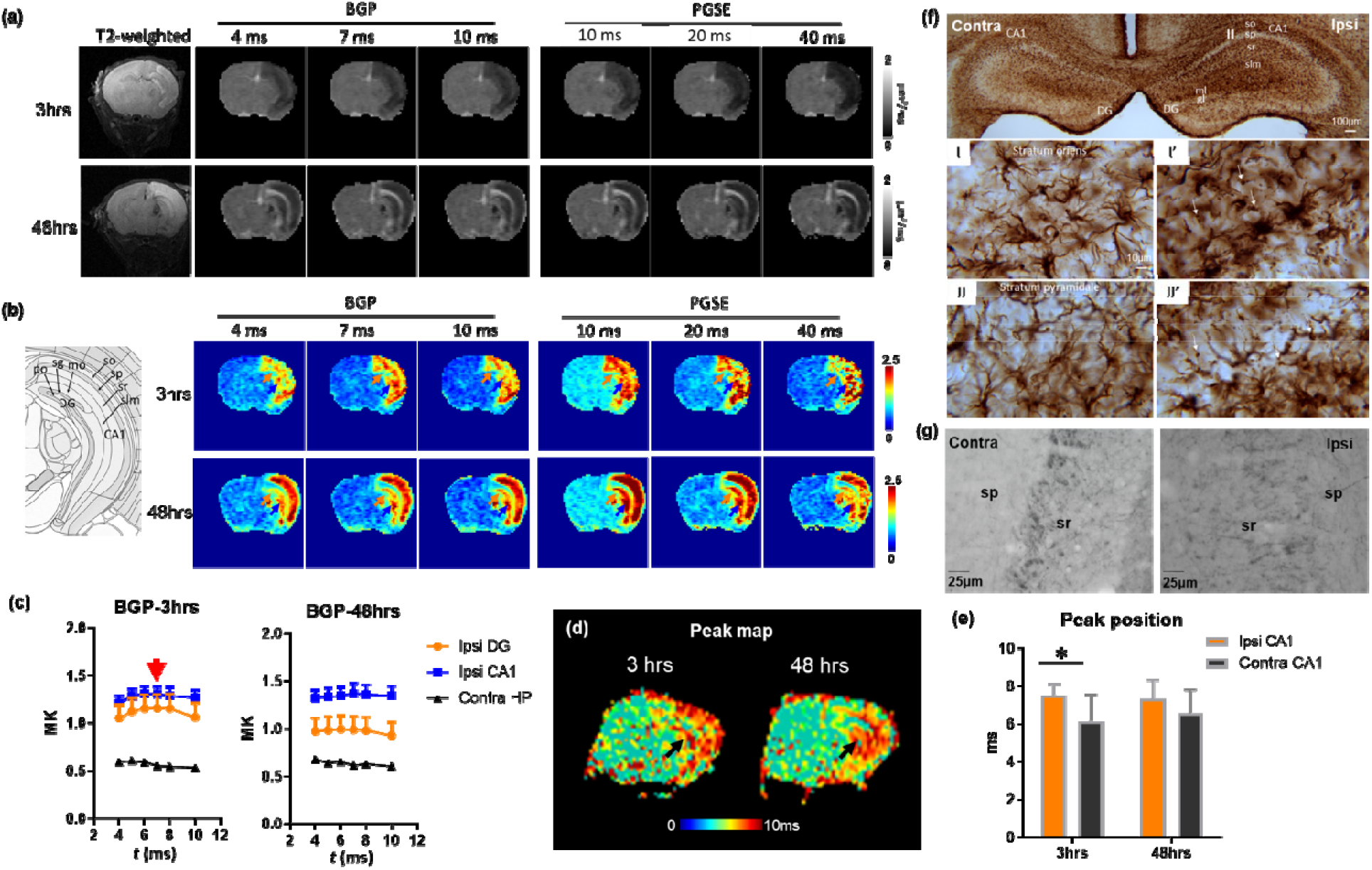
*t*DKI-based *t*_peak_ and microstructural size alterations in a mouse model of neonatal HI injury. (**a-b**) Diffusivity and kurtosis maps of a severely injured mouse brain scanned at 3hrs and 48hrs after the HI onset at *t* from 4-40 ms using bipolar gradient pulse (BGP) and PGSE sequences. (**c**) *t*-dependent change of BGP-based mean kurtosis (MK) showed a peak around 7ms (red arrow) in the ipsilateral CA1, while the contralateral CA1 did not show an apparent peak. (**d**) The *t*_peak_ map showed increased *t*_peak_ in the HP (black arrows) of the HI-injured brain at 3 and 48 hours after HI. (**e**) Statistical comparison of peak positions in the ipsilateral and contralateral CA1 at 3hrs and 48 hrs after HI (*n*=6). Note some of severely injured mice were not imaged longitudinally or suffered from image quality issues. * *p*<0.05 by paired t-test. (**f**) GFAP staining at 3 hours post-HI shows evident beading of astrocytic processes in stratum oriens (I’) and stratum pyramidale (II’) of the ipsilateral hippocampus (thin arrows). (g) Neurofilament staining showed axonal disruption and beading in the ipsilateral stratum pyramidale and stratum radiatum of the ipsilateral hippocampus. Abbreviation: contra—contralateral; ipsi—ipsilateral; sr—stratum radiatum; sp—stratum pyramidale;

In mice that had mild or no apparent edema (about 2/3), slightly decreased diffusivity and increased kurtosis were found in limited regions (mostly in the hippocampus) with T_2_-hyperintensity (**Suppl. Fig S3a,b**). In the group-average kurtosis curves, the peak was not apparent in either ipsilateral hippocampus or the control groups (**Suppl. Fig S3c**).

### tDKI-based permeability indicated tissue survival after HI

We also examined transmembrane water exchange using *t*DKI in the mouse HI model. As shown in **Fig. 5a,b**, no abnormality was visible in the T_2_-weighted images of a severely-injured mice at 3 hrs post injury, but marked changes in diffusivity and kurtosis were already present. Kurtosis maps revealed considerable regional heterogeneity at 3 hrs in the edema region (**Fig. 5b**). In particular, the cingulate cortex (pink arrows) had the highest kurtosis compared to the neighboring regions, e.g. the sensory cortex (light blue arrows), based on which, one might suspect that the cingulate cortex had more severe injury than the sensory cortex, but analyses of the *t*DKI curve indicated otherwise. **Fig. 5c** shows that *t*DKI curve in the sensory cortex descended slightly faster than the cingulate cortex, translating into lower τ*_ex_* (higher permeability) in the sensory cortex (peak arrow in **Fig. 5d**). At 48 hrs, kurtosis in the ipsilateral sensory cortex increased further, and the estimated τ_ex_ was significantly lower than the normal tissue; while the ipsilateral cingulate cortex showed reduced but still elevated kurtosis compared to 3 hrs and τ_ex_ was almost back to normal (**Fig. 5c,e**). Histology at 48 hours demonstrated severe infarct-like necrosis in the ipsilateral sensory cortex, which explained the drastically elevated transmembrane permeability, whereas the cingulate cortex had no apparent infarction and relatively preserved cytology (**Fig. 5f,g** and **Suppl. Fig. S4**). Therefore, the transmembrane permeability marker could be a predictor of tissue viability. The transiently increased permeability in the ipsilateral cingulate at 3hrs, on the other hand, may be explained by the substantial astrocyte activation (**Fig. 5h** and **Suppl. Fig. S5**), which give rise to upregulated water transport^40^ and cytotoxic edema^32,41^.

**Fig. 5:**
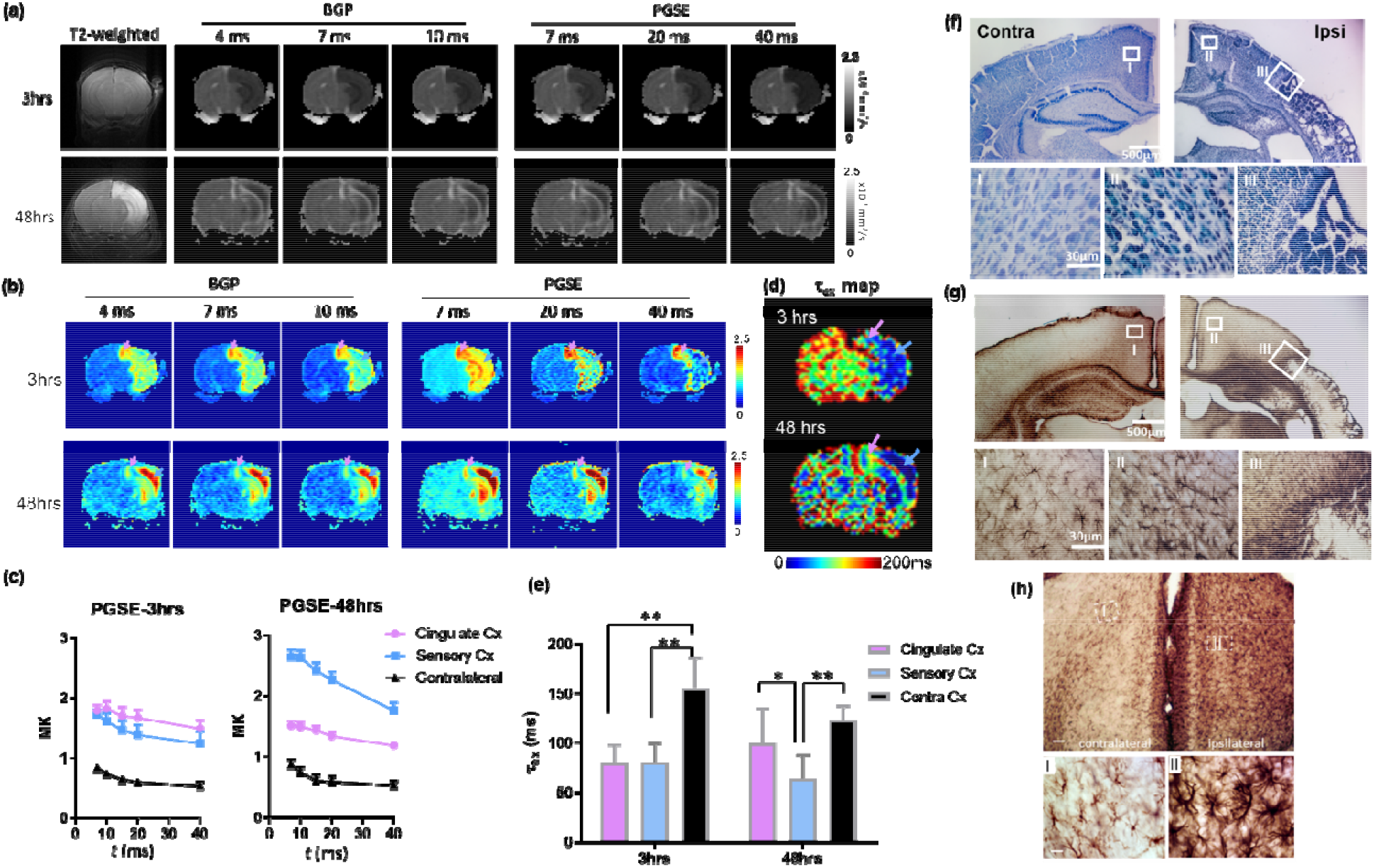
*t*DKI-based estimations of transmembrane water exchange in a mouse model of neonatal HI injury. (**a-b**) Diffusivity and kurtosis maps of a severely injured mouse brain scanned at 3hrs and 48hrs after the HI onset at *t* from 4-40 ms using a combination of bipolar gradient pulse (BGP) and pulsed gradient spin echo (PGSE) sequences. The pink and blue arrows pointed to the cingulate cortex and visual cortex, respectively. (**c**) *t*-dependent change of mean kurtosis (MK) between 7-40 ms illustrated a faster decay tail in the sensory cortex, compared to the nearby cingulate cortex and contralateral side (*n*=6). (**d-e**) At 3 hours after HI, both ipsilateral cingulate (pink arrow) and sensory (blue arrow) cortex had reduced τ*_ex_*, and at 48 hours, τ*_ex_* continued to decrease in the sensory cortex and became lower than that in the cingulate cortex (*n*=6) (* *p*<0.05, ** *p*<0.01 by paired t-test). (**f**) Nissl staining of brain sections at 48 hours after HI reveals a stark difference with the infarcted sensory cortex while the cingulate cortex was almost intact. Higher magnification shows the cellular pathologies in the contralateral (I) and ipsilateral cingulate cortex (II), and the border between ipsilateral cingulate and sensory cortex (III). (**g**) GFAP staining of the same sections as (**f**) shows slight astrocytic activation in the ipsilateral cingulate cortex (II) but much higher astrocyte aggregation on the borders of the necrotic core in the sensory cortex (III). (**h**) GFAP staining of another mouse brain at 3hrs after HI revealed extensive astrocytic activation in the ipsilateral cingulate cortex. High magnification images in **Suppl. Fig. S5** further shows swelling of the astrocytic processes.

In the mild/moderate injury mice, which exhibited reduced diffusivity and increased kurtosis in the hippocampal CA1 region at 3hrs, typically developed apoptotic injury with slightly atrophic hippocampus but no abnormalities in the T_2_-weighted contrast, diffusivity, or kurtosis values at 48 hrs. Quantitative analysis of these mice showed no significant changes in τ*_ex_*, compared to the contralateral side or the shams (**Suppl. Fig. S3d**), indicating no apparent changes in membrane permeability.

### tDKI detected reduced transmembrane exchange in an ischemic stroke patient

Encouraged by the findings from the mouse HI model, we went one step further to see clinical feasibility of *t*DKI in ischemic stroke. A stroke patient (male, 66 years old) with a right posterior cerebral artery infarction was scanned at approximately 36 hours after onset, on a 3T Siemens scanner using STEAM diffusion encoding with *t* from 50-250ms. Note that given the vast difference in the size of cell body and processes between mouse and human brains ^42^, we choose a long *t* range to ensure the completion of coarse graining in the human study. **Fig. 6a-b** demonstrated elevated kurtosis and reduced diffusivity in the injured tissues. Kärger model fitting revealed decreased exchange time τ*_ex_* and increased *K_0_* (kurtosis at infinitely short *t*) in the edematous tissue (thick arrow), which agreed with the reduced τ*_ex_* in the HI-injured mouse brains. Regional heterogeneity with high τ*_ex_* along the posterior white matter (thin arrow) may related to the intrinsically low permeability in the white matter due to the myelin sheath ^28^, and the high *K_0_* could indicate restricted diffusion due to white matter edema ^9^.

**Fig. 6:**
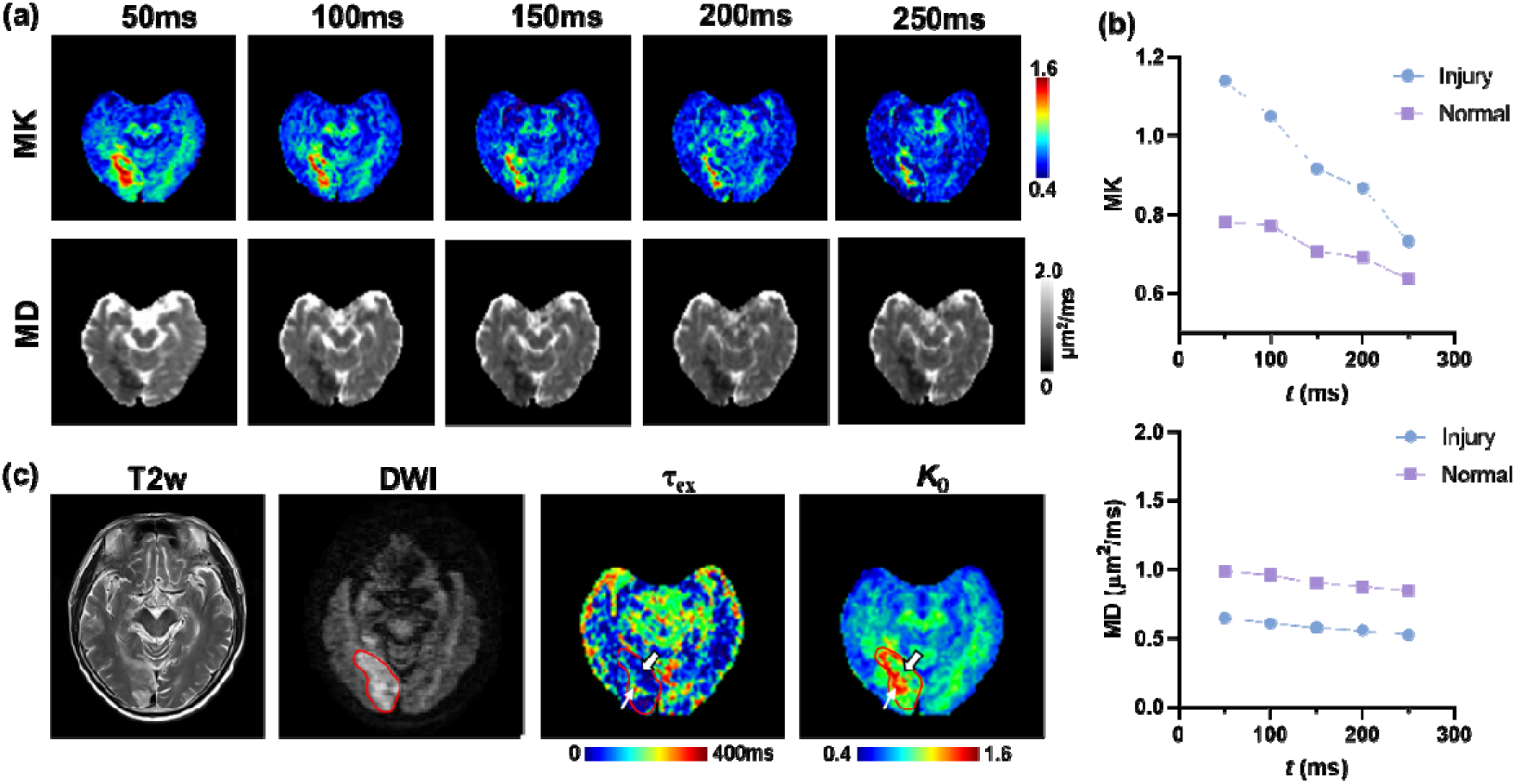
tDKI in an ischemia stroke patient. (**a**) The mean kurtosis (MK) and mean diffusivity (MD) maps at each *t* from 50 to 250 ms. (**b**) Diffusion-time dependent MK and MD curves in the edema tissue (red contour, partly delineating the optic radiation) and the normal tissue in the contralateral side. (**c**) The T2-weighted image, diffusion-weighted image (DWI), and exchange time τ_ex_ and intrinsic *K_0_* as estimated from the Kärger model. Thick arrows point to reduced τ_ex_ and increased *K_0_* in the injured brain tissues, while thin arrows to the white matter within the edema region.

## Discussion

The *t*DKI provided a non-invasive real-time approach to detect the pathological events after ischemic injury at the cellular level, well beyond the resolution of conventional MRI. The pathophysiology of cerebral ischemia involves a chain of complex cellular and molecular processes, and *in vivo* mapping of such events is essential to identify the tissue damage and viability and design therapeutic strategies. During the process of HI, ionic and osmotic perturbations occur resulting in water shifting to intracellular compartments as well as swelling and beading of cell body and processes. The timing of neuropathology within different microdomains of neural cells is unclear and difficult to identify using conventional dMRI. Our result indicates that the morphological change detected by peak of the *t*DKI curve is likely associated with astrocytic processes and neurites. Simultaneously to the morphological changes, alternations in membrane permeability are also measured using tail of *t*DKI curve, not only reflecting the transiently upregulated astrocytic function but also disrupted membrane integrity that is vital to tissue outcome.

The observed *t*-dependent behaviors of diffusivity and kurtosis measurements hinted to the spatial scale of structural barriers. The relatively short *t*_peak_ of the *t*DKI curve (≤10 ms) from *in vitro* and *in vivo* experiments suggests that the *t*DKI measurement is primarily associated with structural barriers at the level of axons, dendrites, or glial processes, instead of cell body. As the *t*DKI peak of normal tissue appeared to be shorter than 4 ms, the peak at 7-8 ms in acute edema regions might reflect the observed swelling/beading of the astrocytic processes or neurites (axons and dendrites) with comparable sizes (∼2 µm). Histopathology in this study show more widespread swelling of the astrocyte compartments compared to sporadic axon beadings, suggesting that astrocytes may play a major role here, although the definite contribution from the two sources remain difficult to quantify. Previous studies showed that the astrocytes play a critical regulatory effect in ischemic stroke ^43^ and have become a potential treatment target ^44^. For instance, astrocytes rapidly sense the ischemic insult and rapidly tune their volumes and regulate their processes ^32^, partially by upregulating the AQP4 channels on its endfeets to facilitate transmembrane water exchange ^40,45^. The *t*DKI findings of microstructural size and permeability changes during the early evolving cytopathology of brain HI is consistent with predominance of protoplasmic astrocytes and their processes represented in the neuropil ^46^, and therefore, provide the possibility for *in vivo* detection of astrocytic pathology.

Inspecting membrane integrity via water exchange using *t*DKI requires the completion of the coarse-graining. When the restrictive effect of microstructure is accounted, the *t*-dependent change in kurtosis mainly reflects exchange effects, and a faster decay of the *t*DKI tail corresponds to higher permeability (Fig. 1). Our mouse model suggests that *in vivo* mapping of the transmembrane exchange or permeability correlates with the tissue integrity outcome. For example, at 3hrs after HI, the lower kurtosis in the ipsilateral sensory cortex than cingulate cortex at single diffusion-times was misleading, and contradictory to the final outcome. *t*DKI revealed higher permeability in the sensory cortex that matched to its final progression into necrosis and severe tissue infarction. This is promising in identification and prediction of salvageable tissues for which therapies can be useful. We further demonstrated that this technique was applicable in ischemic stroke patient, who showed increased permeability in injured brain regions at subacute stage after reperfusion, a time when irreversible injury occurred.

A variety of dMRI techniques have been proposed for neurite diameter mapping, including *q*-space approaches by acquiring dMRI signals in multiple diffusion directions and strengths ^47–49^ as well as *t*-dependent dMRI methods ^16,50^. Most of these approaches involved complex multi-compartmental modeling and lengthy sampling in the *q*-space or *t*-domain. On the other side, methods for spatial mapping of transmembrane permeability are relatively few. The filter-exchange imaging (FEXI) method ^51^ is thought to infer the exchange time by modulating the intracellular (low diffusivity) and extracellular (high diffusivity) signals with a variable exchange time, which has been tested in animal models ^52^ and human brain ^53^ but the signal-to-noise ratio is of concern. The random permeable barrier model also allows for permeability estimation based on the power-law of time-dependent diffusion ^13,14,25,54,55^. Here, we demonstrated the use of *t*DKI for simultaneously assessing structural morophology and permeability based on the *t*DKI curve and illustrated the clinical value of these microstructural markers in ischemic injury. The clinical version of *t*DKI technique only takes 10 minutes, and thus can be readily translated to clinical applications.

To take the full advantage of *t*DKI, it is ideal to have a wide range of *t*s from ∼1ms to ∼100ms, which is challenging using the same type of diffusion encoding. We used the BPG ^56^ encoding to access *t* <10 ms, PGSE ^57^ diffusion encoding for 10ms < *t* < 50 ms, and STEAM sequence ^58^ for *t* > 50 ms. During calibration, we found that PGSE and STEAM provided consistent kurtosis measurement, while BPG exhibited a bias (**Suppl. Fig S6**). Therefore, in the current study, we combined PGSE and STEAM measurements in the *in vitro* organoid experiment; while in the mouse brain study, we separately used BPG data to estimate the *t*DKI peak and PGSE data to fit the tail. When it comes to clinical applications, measurements at short *t* become difficult given the limited gradient strength (only ∼1/10 of the animal scanners). Instead, we focused on the *t*DKI tail at long *t* (50-250 ms) to map transmembrane permeability using the STEAM sequence.

This study has several limitations. First, only a selected range of *t’*s is acquired with each diffusion encoding scheme due to the technical constraints discussed above and practical considerations, as sampling a large number of *t*s is not feasible for *in vivo* studies. For future studies and potential clinical applications, further optimization of the set of *t*s is needed. Second, *t*DKI curve reflected a mixed effect of microstructural size and permeability, which requires complete coarse graining to separate these two effects. Multi-dimensional MRI combining relaxometry and diffusion ^59^ may help to further disentangle the two effects. Furthermore, we did not directly measure the transmembrane permeability in this study, but inferred the permeability change from histological observations of astrocyte physiopathology. Future studies on mice expressing fluorescent proteins completely in astrocytes to visualize their entire volumes and aquaporin or Na/K-ATPase antibodies could provide more direct evidence of transmembrane water exchange. In addition, the fitted τ*_ex_* maps were relatively noisy given the Kärger model fitting and limited data points. Bayesian fitting or machine learning approaches ^60^ may be attempted to improve the fitting results.

Nevertheless, the novel findings presented here using *t*DKI provided important information on structural morphology and transmembrane permeability in addition to existing MRI routines, and a tentative suggestion of its potential to detect astrocytic pathology and predict pathological outcomes related to reversible and salvageable or irreversible damage in ischemic stroke. Further applications of this technique in other diseases are also promising, such as tumor where considerable changes in cellular microstructure and permeability take place.

## Acknowledgments

This work is supported by Ministry of Science and Technology of the People’s Republic of China (2021ZD0200202, D.W.), National Natural Science Foundation of China (81971606 and 82122032, D.W.), Science and Technology Department of Zhejiang Province (202006140 and 2022C03057, D.W.), National Institutes of Health (R01NS102904 and R01HD074593, J.Z., NS114144 / NS123814 / HD110091 / HD074593, F.J.N., NS079348 and HD074593, L.J.M.), and Johns Hopkins University Alzheimer’s Disease Research Center (AG061643, L.J.M), and was performed at the Center of Advanced Imaging Innovation and Research (CAI2R, www.cai2r.net), a Biomedical Technology Resource Center supported by National Institute of Biomedical Imaging and Bioengineering of the NIH under the award P41 EB017183.

## Materials and Methods

### Monte Carlo simulation

We performed Monte Carlo simulation ^33,34^ for cylinder and sphere phantoms to represent simplified geometry of axons and cells, respectively. The neurite phantoms were configured in parallel to each other with random packing at diameters of 1-1.4 µm, varying transmembrane exchange times of 10-200 ms, intra-neurite volume fraction of 0.3 for 1 µm which increased with the neurite diameter, and fixed intra- and extra-neurite diffusivities of 1 and 2 µm^2^/ms, respectively. The cell body phantoms were configured similarly, with the diameters set at 10-14 µm. dMRI signals were sampled at 1000 *t*’s from 0.2-200 ms, b-values from 0.5-3 ms/ µm^2^, and 10 diffusion directions (radial to the neurite orientation and isotropic for cell) per b-value. Diffusivity and kurtosis were calculated using the simulated signals, according to 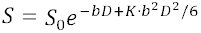, for each of the microstructural configurations and the diffusion encoding schemes.

### Diffusion MRI pulse sequence

To access water diffusion experimentally over a wide range of *t*, three types of diffusion gradient waveforms were employed. We used the bipolar pulsed gradient (BPG) ^56^ to measure dMRI signals at short *t*’s (3-10ms), the pulsed gradient spin-echo (PGSE) ^57^ for intermediate *t*’s (7-40ms), and the stimulated echo acquisition mode (STEAM) ^58,61^ for intermediate and long *t*’s (15-200ms). The sequence schematics and their effective *t*’s were defined in Suppl Fig. S1. These sequences were calibrated using a mineral oil phantom to ensure consistent diffusivity measurements.

Note that kurtosis measurements in biological tissue may differ among the different diffusion encoding schemes, even at comparable *t*, as reported previously^62^. In our experimental data (Suppl Fig. S5), we found BPG and PGSE resulted in similar diffusivity but different kurtosis at the same *t*, while PGSE and STEAM provided consistent results. Therefore, we analyzed the tDKI curves obtained by BPG and STEAM separately in the *in vivo* mouse brain study, while used the PGSE and STEAM measurements jointly in the organoid experiment.

### Forebrain organoid and in vitro MRI

The 3D brain organoids (3Dnamics, Inc., Baltimore, MD, USA) differentiated from human induced pluripotent stem cells (hiPSCs) to resemble the molecular and cellular features of the human forebrain. Th hiPSCs derived from human adult somatic cells were cultured in forebrain differentiation medium, according to the protocol described in ^63^. The 3D organoids were placed in glass tubes surrounded agarose gel for supporting and preventing hydration during scan. They were scanned on an 11.7T Bruker vertical spectrometer, with a 15 mm volume transceiver coil, a Micro2.5 gradient system (maximum gradient strength = 1500 mT/m), and a temperature control system. dMRI data were acquired using PGSE sequence with diffusion gradient duration (δ) of 3 ms, diffusion separation (Δ) of 7-30 ms (equivalent *t* = Δ-δ/3), and two signal average; and STEAM sequence with δ of 3 ms, Δ of 15-200 ms, and eight signal average to compensate for the SNR loss in STEAM. The other diffusion parameters were kept the same between PGSE and STEAM, including 30 diffusion directions, six b-values with b = 0.5, 1.0, 1.5, 2.0, 2.5, and 3.0 s/mm^2^, and five non-diffusion-weighted images. The effective b-values that considered the contributions of imaging gradients were calculated on the Bruker system and used in the following analyses. dMRI images were acquired using a single-shot 3D echo planar imaging (EPI) and the following parameters: echo time (TE) / repetition time (TR) = 38/3000 ms, in-plane resolution = 0.31 mm × 0.31 mm × 0.8 mm, and an imaging time of 18.5 minutes for PGSE and 1.23 hours for STEAM scans. The organoids were first imaged room temperature (300K), and then the scans were repeated at 305K, 310K, 315K, and 320K. We left at least 3 hours interval for the temperature transitions.

### Mouse model of neonatal hypoxia-ischemia

All animal procedures were approved by the Animal Use and Care Committee of Johns Hopkins University School of Medicine. In this study, 28 C57BL/6 mouse pups (Jackson Laboratory, Bar Harbor, ME, USA) underwent HI insult on postnatal day 10 (P10), using the Rice-Vannucci model adapted for the neonatal mouse ^64^. The injury was induced by unilateral ligation of the right carotid artery followed by 45 minutes of hypoxia (FiO2=0.08), as described in ^64^. 15 pups (1-2 from each litter) were subjected to sham injury, with right carotid artery exposed but no ligation or hypoxia. The first MRI was acquired at 3-6 hours after the hypoxia. Some of the mice were killed after imaging for pathology, and the rest of the them were survived up to 48 hours until they were scanned again and then killed for pathology. The use of the mice is specified in Suppl. Table S3.

### In vivo MRI of the mouse brains

*In vivo* MRI was performed on a horizontal 11.7 Tesla scanner (Bruker Biospin, Billerica, MA, USA). MR images were acquired with a 15 mm receive-only planner surface coil, a 72 mm quadrature transmitter coil, and a B-GA 9S gradient system (maximum gradient strength = 740 mT/m). During imaging, mice were anesthetized with isoflurane (1%) together with air and oxygen mixed at 3:1 ratio via a vaporizer. PGSE sequence was acquired at δ = 4 ms and Δ = 7, 10, 15, 20, 40 ms, and BPG sequence at δ = 3 ms and Δ = 4.1, 5, 6, 7, 8, 10 ms. All dMRI protocols used 15 diffusion directions, three b-values with b = 1.0, 1.5, 2.0 ms/µm^2^, four non-diffusion-weighted images. Both the PGSE and BPG scans were acquired using single-shot 2D multi-slice EPI with TE/TR = 55/5000 ms, one signal average, in-plane resolution = 0.2 mm × 0.2 mm, 10 slices with slice thickness of 0.8 mm, and an imaging time of 4.1 minutes per scan. In addition, T2-weighted images were acquired using a fast spin echo sequence with TE/TR = 50/3000 ms, two signal averages, echo train length of 8, in-plane resolution of 0.1 mm x 0.1 mm, and 10 slices with thickness of 0.80 mm, co-registered to the dMRI data.

### Histology and Immunohistochemistry for neuropathology

Mice were perfused intracardially with phosphate-buffered saline followed by 4% paraformaldehyde. Dissected brains were cyroprotected in sucrose and sectioned at a thickness of 50 µm on a freezing sliding-microtome. Every 10^th^ section was stained with cresyl violet to visualize Nissl substance and adjacent sections were stained with hematoxylin and eosin (H&E). Near-adjacent sections were stained immunohistochemically using the peroxidase anti-peroxidase method for glial fibrillary and acidic protein (GFAP) as an astrocyte marker, MAP2 for a neuron cell body and dendrite marker, and a phosphorylated neurofilament antibody as a marker for axonal processes, and microtubule protein MAP2 for dendrites.

### tDKI acquisition of ischemic stroke patients

An ischemic stroke patient (male, 66 years old) was assessed in this study, who underwent thrombolysis at 3 hours after onset. The study was approved by the Institutional Research Board at Weifang Medical School and written consent was collected from the patient. tDKI scan was performed at 4 days after stroke on a 3T Siemens Skyra scanner (maximum gradient strength = 45 mT/m) using the STEAM diffusion MRI sequence with the following protocol: TE/TR = 53/2500 ms, FOV = 220 × 220 ms, resolution = 2.3 × 2.3 ms, 14 slices with slice thickness of 5 mm, 20 diffusion directions per b-value at two b-values of 1 and 2 ms/µm^2^, and effective *t*s of 50, 100, 150, 200, and 250 ms. Total scan time was about 10 mins.

### Data processing

Mean diffusivity and mean kurtosis at each *t* were estimated using the matlab routine developed by Dr. Jelle Veraart (https://github.com/jelleveraart/RobustDKIFitting). Based on the tDKI curves, we fitted the peak positions using a Gaussian model, in which the center of the Gaussian corresponded to the peaks position. For those curves that did not show a biphasic shape, the peak position was set to the shortest *t* in measurement (lower limit of the Gaussian center). The transmembrane exchange time (τ*_ex_*) were fitting based on the Kräger model ^35^:

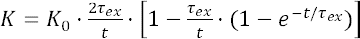

where *K_0_* is the kurtosis at infinitely short diffusion time and τ*_ex_* is the exchange time (dwell time of water molecules within one compartment), which is inversely related to exchange rate.

Regions of interest (ROIs) were manually defined, e.g., the rim and core of the organoids, and the cingulate cortex, occipital cortex, dentate gyrus (DG) and CA1/CA3 regions of the hippocampus in the mouse brains. The ROI-averaged diffusivity and kurtosis values were obtained for statistical analysis.

### Statistical analysis

In the organoid experiment, the fitted kurtosis peak positions and τ*_ex_* were compared between the rim and core and between different temperatures, using two-way analysis of variance (ANOVA), followed by posthoc *t*-test with Sidak correction for multiple comparisons in Graphpad Prism (http://www.graphpad.com/scientific-software/prism/).

In the *in vivo* mouse brain experiment, the diffusivity and kurtosis were compared between different ROIs and between two time points at individual *t*, using multi-way ANOVA, followed by posthoc pairwise *t*-test with Sidak correction for multiple comparisons. The statistical differences were listed in Suppl. Table S4-6. The fitted τ*_ex_* or peak positions were compared between different ROIs and between 3hrs and 48hrs, using two-way ANOVA, followed by posthoc *t*-test with Sidak correction.

## Notes

### Competing Interest Statement

The authors have declared no competing interest.

