## supplementary materials for "*In vivo* Mapping of Cellular Resolution Neuropathology in Brain Ischemia by Diffusion MRI"

**Supplementary Tables**

**Supplementary Table S1:** Peak position of *t*DKI curve estimated from simulated dMRI signals based on varying diameter sand transmembrane water exchange times in the neurite and cell phantoms.

| Peak posotion (ms) | | τ*_ex_* (ms) used in simulations | | | | | | | | | | | | |
| --- | --- | --- | --- | --- | --- | --- | --- | --- | --- | --- | --- | --- | --- | --- |
|  |  | 10 | 20 | | 30 | | 40 | | 50 | | 100 | | 200 | |
| **Neurite**  *d* (µm) | 1.0 | 1.0 | | 1.4 | | 1.4 | | 1.8 | | 1.6 | | 1.6 | | 2.2 |
|  | 1.1 | 1.6 | | 1.8 | | 1.8 | | 2.4 | | 2.0 | | 2.0 | | 2.8 |
|  | 1.2 | 1.6 | | 2.2 | | 2.6 | | 2.6 | | 3.2 | | 3.4 | | 3.4 |
|  | 1.3 | 1.8 | | 2.2 | | 2.6 | | 3.2 | | 3.4 | | 3.8 | | 5.8 |
|  | 1.4 | 2.2 | | 3.0 | | 3.4 | | 4.0 | | 4.2 | | 5.2 | | 5.8 |
| **Cell**  *d* (µm) | 10 | 11.0 | | 19.8 | | 22.8 | | 29.6 | | 34.8 | | 47.2 | | 69.4 |
|  | 11 | 12.4 | | 22.2 | | 28.0 | | 31.0 | | 38.4 | | 56.0 | | 69.2 |
|  | 12 | 10.4 | | 25.6 | | 32.4 | | 40.2 | | 40.6 | | 61.8 | | 96.2 |
|  | 13 | 11.4 | | 28.6 | | 37.8 | | 45.6 | | 51.8 | | 71.8 | | 99.4 |
|  | 14 | 14.2 | | 34.6 | | 44.0 | | 48.8 | | 57.6 | | 89.8 | | 121.0 |

**Supplementary Table S2:** Estimated transmembrane exchange time (τ*_ex_*) of *t*DKI curve estimated from simulated dMRI signals based on varying diameters and τ*_ex_* in the neurite and sphere phantoms.

| Estimated τ*_ex_* (ms) | | τ*_ex_* (ms) used in simulations | | | | | | | | | | | | |
| --- | --- | --- | --- | --- | --- | --- | --- | --- | --- | --- | --- | --- | --- | --- |
|  |  | 10 | 20 | | 30 | | 40 | | 50 | | 100 | | 200 | |
| **Neurite**  *d* (µm) | 1.0 | 9.7 | | 19.6 | | 29.2 | | 38.1 | | 48.3 | | 97.7 | | 185.5 |
|  | 1.1 | 9.7 | | 19.5 | | 29.1 | | 38.6 | | 49.2 | | 94.1 | | 186.1 |
|  | 1.2 | 9.9 | | 19.4 | | 29.7 | | 38.8 | | 48.4 | | 96.3 | | 187.7 |
|  | 1.3 | 10.2 | | 19.6 | | 29.3 | | 39.1 | | 49.1 | | 96.7 | | 187.6 |
|  | 1.4 | 10.3 | | 20.2 | | 29.7 | | 39.3 | | 49.5 | | 96.7 | | 192.4 |
| **Cell**  *d* (µm) | 10 | 17.2 | | 29.1 | | 42.6 | | 55.3 | | 66.0 | | 135.2 | | 282.2 |
|  | 11 | 18.7 | | 32.1 | | 45.2 | | 59.4 | | 72.0 | | 138.1 | | 358.5 |
|  | 12 | 21.5 | | 34.3 | | 47.8 | | 62.3 | | 81.4 | | 164.4 | | 341.0 |
|  | 13 | 23.5 | | 37.0 | | 53.0 | | 66.8 | | 82.2 | | 174.7 | | 454.3 |
|  | 14 | 25.6 | | 41.6 | | 56.7 | | 77.6 | | 95.7 | | 199.0 | | 620.9 |

**Supplementary Table S3:** Summary of the *in vivo* mouse brain experiment of neonatal HI injury.

|  | Severe injury | Mild/moderate | Sham |
| --- | --- | --- | --- |
| 3hr MRI | 12 | 10 | 6 |
| 48hr MRI | 8 (follow-up) | 5 (follow-up) | 6 |

**Supplementary Table S4:** Statistical analysis associated with Fig. 4c, differences in mean diffusivity (MD) and mean kurtosis (MK) between the ipsilateral and contralateral hippocampus of the severely HIE mice (*n*=12), as well as the sham mice (*n*=6), at 3 hrs after HI injury. The comparisons were performed at individual *t* from 4-10 ms using the bipolar pulsed gradient (BPG) and from 7-40 ms using the pulse gradient spin-echo (PGSE) sequence. The differences between each pair of comparison were shown. * *p*<0.05, ** *p*<0.01, and *** *p*<0.001 by t-test.

|  | MD | | | MK | | |
| --- | --- | --- | --- | --- | --- | --- |
|  | Ipsi - Contra | Ipsi - Sham | Contra - Sham | Ipsi - Contra | Ipsi - Sham | Contra - Sham |
| BPG 4 ms | -0.046** | -0.020 | 0.025 | 0.136*** | 0.092*** | -0.044 |
| BPG 5 ms | -0.049** | -0.026 | 0.023 | 0.145*** | 0.111*** | -0.033 |
| BPG 6 ms | -0.044** | -0.036* | 0.008 | 0.152*** | 0.088*** | -0.065* |
| BPG 7 ms | -0.053*** | -0.026 | 0.027 | 0.125*** | 0.106*** | -0.019 |
| BPG 8 ms | -0.048*** | -0.027 | 0.021 | 0.147** | 0.108** | -0.039 |
| BPG 10 ms | -0.059*** | -0.037** | 0.022 | 0.076 | 0.070 | -0.006 |
| PGSE 7 ms | -0.034 | -0.005 | 0.029 | 0.137* | 0.101 | -0.036 |
| PGSE 10 ms | -0.034 | -0.018 | 0.016 | 0.156** | 0.111 | -0.045 |
| PGSE 15 ms | -0.044 | -0.030 | 0.014 | 0.145* | 0.120* | -0.025 |
| PGSE 20 ms | -0.061 | -0.029 | 0.032 | 0.085 | 0.087 | 0.002 |
| PGSE 40 ms | -0.056 | -0.037 | 0.019 | 0.102 | 0.080 | -0.022 |

**Supplementary Table S5:** Statistical analysis associated with Fig. 5c, comparing MD values in the cingulate and sensory cortex of the HIE mice at 3 hrs (*n*=12) and 48 hrs (*n*=8) after HI injury, at *t* from 4-40ms. The differences between each pair of comparison were shown. * *p*<0.05, ** *p*<0.01, and *** *p*<0.001 by paired t-test.

| MD | 3 hrs | 48 hrs | | | 48hrs – 3hrs | |
| --- | --- | --- | --- | --- | --- | --- |
|  | Cingulate - Sensory | Cingulate - Sensory | Cingulate – contralateral | Contra - contralateral | Cingulate | Sensory |
| BPG 4 ms | 0.0296 | 0.2505*** | 0.0339 | -0.2166*** | -0.1608*** | 0.0601 |
| BPG 5 ms | 0.0460 | 0.2407*** | 0.0326 | -0.2081*** | -0.1689*** | 0.0258 |
| BPG 6 ms | 0.0634 | 0.2554*** | 0.0289 | -0.2265*** | -0.1787*** | 0.0133 |
| BPG 7 ms | 0.0600 | 0.2526*** | 0.0334 | -0.2192*** | -0.1809*** | 0.0118*** |
| BPG 8 ms | 0.0628 | 0.2620*** | 0.0344 | -0.2276*** | -0.1866*** | 0.0126 |
| BPG 10 ms | 0.0651 | 0.2688*** | 0.0385 | -0.2304*** | -0.2047*** | -0.0010 |
| PGSE 7 ms | 0.0432 | 0.2544*** | -0.005 | -0.2595*** | -0.1524** | 0.0588 |
| PGSE 10 ms | 0.0293 | 0.2697*** | 0.0114 | -0.2583*** | -0.1757*** | 0.0647 |
| PGSE 15 ms | 0.0301 | 0.2807*** | 0.0291 | -0.2516*** | -0.1969*** | 0.0537 |
| PGSE 20 ms | 0.0364 | 0.2785*** | 0.0305 | -0.2480*** | -0.2119*** | 0.0302 |
| PGSE 40 ms | 0.0385 | 0.2789*** | 0.0261 | -0.2528*** | -0.2435*** | -0.0031 |

**Supplementary Table S6:** Statistical analysis associated with Fig. 5c, comparing MK values in the cingulate and sensory cortex of the HIE mice at 3hrs (*n*=12) and 48hrs (*n*=8) after HI injury, at *t_d_* from 4-40ms. The differences between each pair of comparison were showed. * *p*<0.05, ** *p*<0.01, and *** *p*<0.001 by paired t-test.

| MK | 3 hrs | 48 hrs | | | 48hrs – 3hrs | |
| --- | --- | --- | --- | --- | --- | --- |
|  | Cingulate - sensory | Cingulate - sensory | Cingulate – contralateral | Contra - contralateral | Cingulate | sensory |
| BPG 4 ms | 0.1139 | -0.8904*** | 0.4999** | 1.390*** | 0.3239 | -0.6804*** |
| BPG 5 ms | 0.1435 | -0.9258*** | 0.5647** | 1.490*** | 0.3935 | -0.6757*** |
| BPG 6 ms | 0.1213 | -0.9200*** | 0.5815*** | 1.502*** | 0.3753 | -0.6660*** |
| BPG 7 ms | 0.1155 | -0.9219*** | 0.6148*** | 1.537*** | 0.4408* | -0.5965*** |
| BPG 8 ms | 0.1269 | -0.9460*** | 0.5634** | 1.509*** | 0.4642* | -0.6087*** |
| BPG 10 ms | 0.1924 | -0.9548*** | 0.5728** | 1.528*** | 0.4599* | -0.6873*** |
| PGSE 7 ms | 0.0602 | -1.130*** | 0.6384*** | 1.769*** | 0.2751 | -0.9156*** |
| PGSE 10 ms | 0.2055 | -1.137*** | 0.7467*** | 1.884*** | 0.3259 | -1.017*** |
| PGSE 15 ms | 0.2459 | -1.016*** | 0.7816*** | 1.797*** | 0.3073 | -0.9545*** |
| PGSE 20 ms | 0.2891 | -0.9254*** | 0.7476*** | 1.673*** | 0.3338 | -0.8808*** |
| PGSE 40 ms | 0.2469 | -0.5768** | 0.6544*** | 1.231*** | 0.3043 | -0.5194* |

**Supplementary Figures**

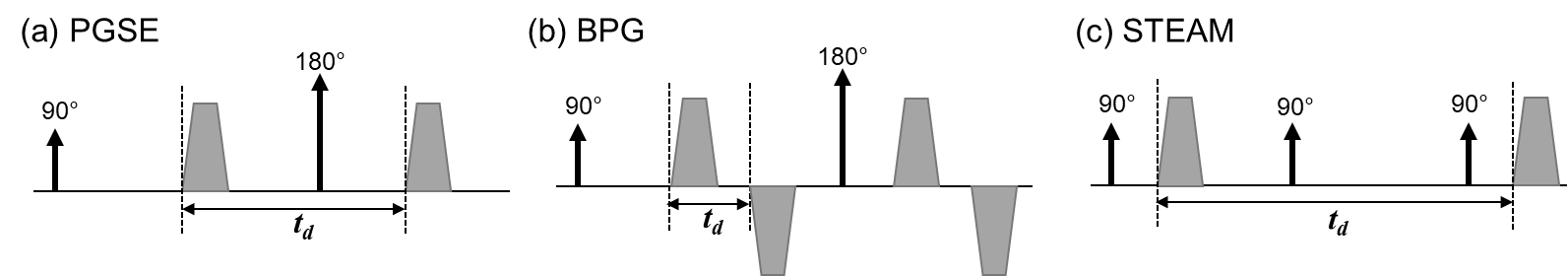

**Fig. S1:** Schematics of the pulse sequences for different diffusion encoding schemes, including (**a**) pulsed gradient spin-echo (PGSE), (**b**) bipolar pulsed gradient (BGP), and (**c**) stimulated echo acquisition mode (STEAM). The effective diffusion time (*t_d_*) are indicated in each scheme.

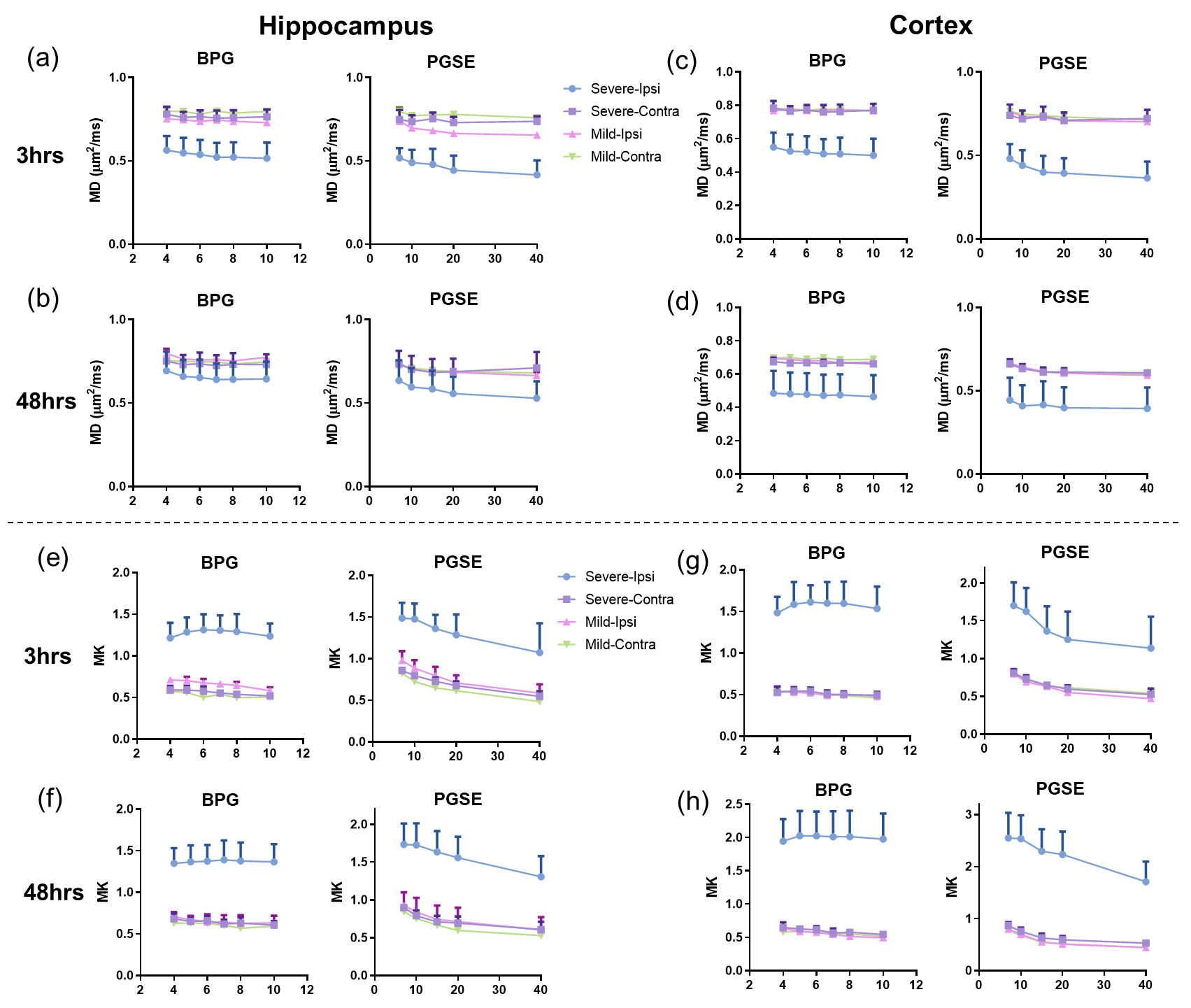

**Fig. S2:** *t*-dependent change of mean diffusivity (MD, **a-d**) and mean kurtosis (MK, **e-h**) measured in the hippocampus and cortex of HI-injured mouse brains at 3hrs and 48hrs after injury, using BGP and PGSE sequences.

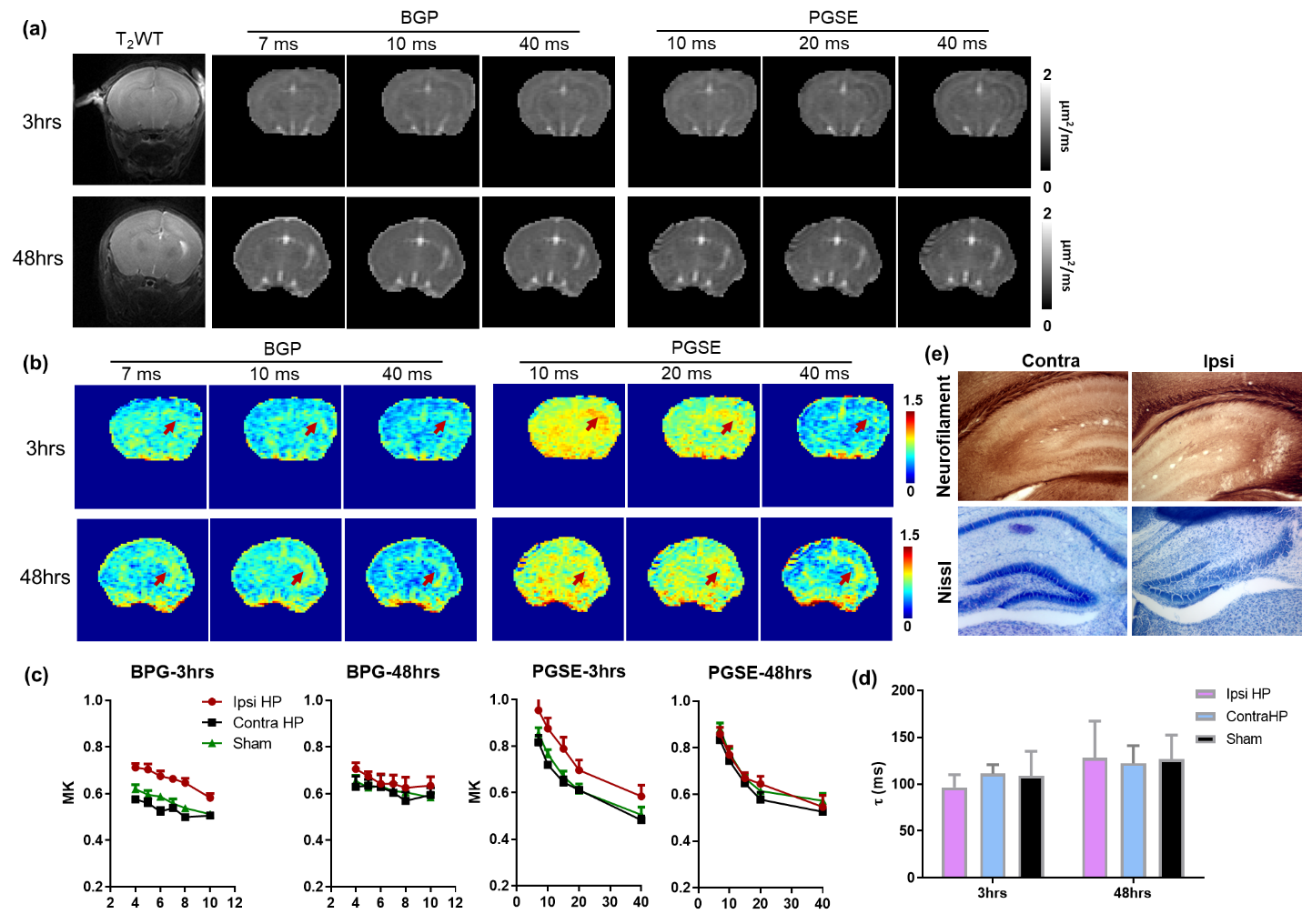

**Fig. S3:** tDKI in HI-injured mice with mild-to-moderate injury. (**a-b**) Diffusivity and kurtosis maps of a severely injured mouse brain scanned at 3hrs and 48hrs after the HIE onset at *t_d_* from 4-40 ms using a combination of BPG and PGSE sequences. The red arrows pointed to elevated kurtosis in the ipsilateral hippocampus. (**c**) *t*-dependent change of mean kurtosis (MK) between 4-10 ms did not show apparent kurtosis peak, and kurtosis tail showed similar rate of decay between 7-40 ms. (**d**) Statistical comparison of τ_ex_ between the ipsilateral and contralateral hippocampus and sham mice did not show statistical difference, either at 3hrs (n=10) or 48 hrs (n=5) after HIE. (**e**) Neurofilament and Nissl staining of the mouse brain at 48 hrs after HIE showed largely intact hippocampus without apparent axonal swelling or neuronal damage.

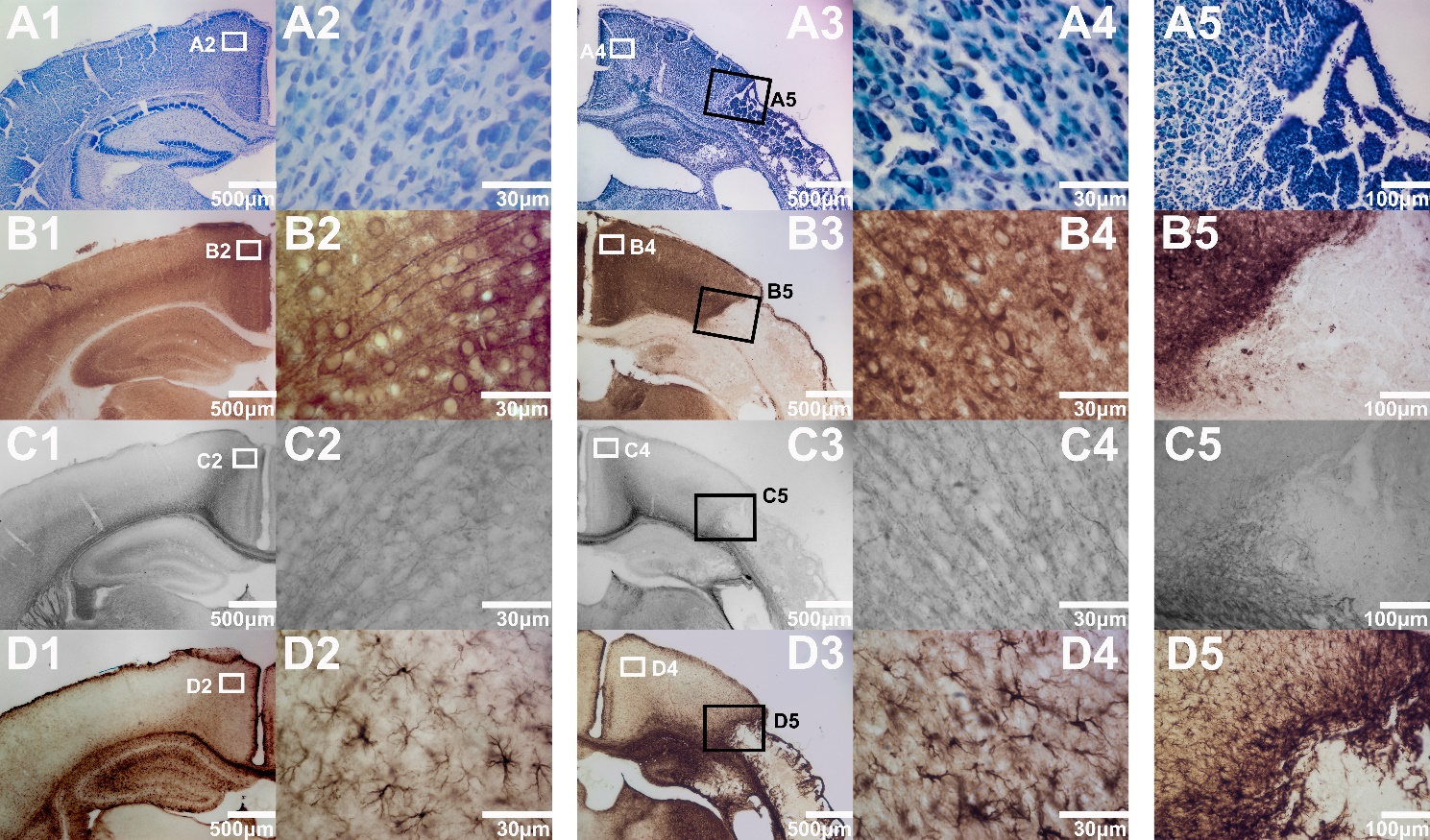

**Fig. S4:** Pathology of the neurons, astrocytes, and cellular processes of a neonatal brain at 48 hours after HI injury, corresponding to the mouse brain in Fig. 5. (A) Nissl staining of the contralateral (A1) and ipsilateral (A3) brain with zoom-in views of the cingulate cortex (A2 and A4) and the junction between cingulate and sensory cortex (A5). Same configuration is used for MAP2 (B), Neurofilament (C), and GFAP (D) to characterize the pathology of the dendrite, axons, and astrocytes.

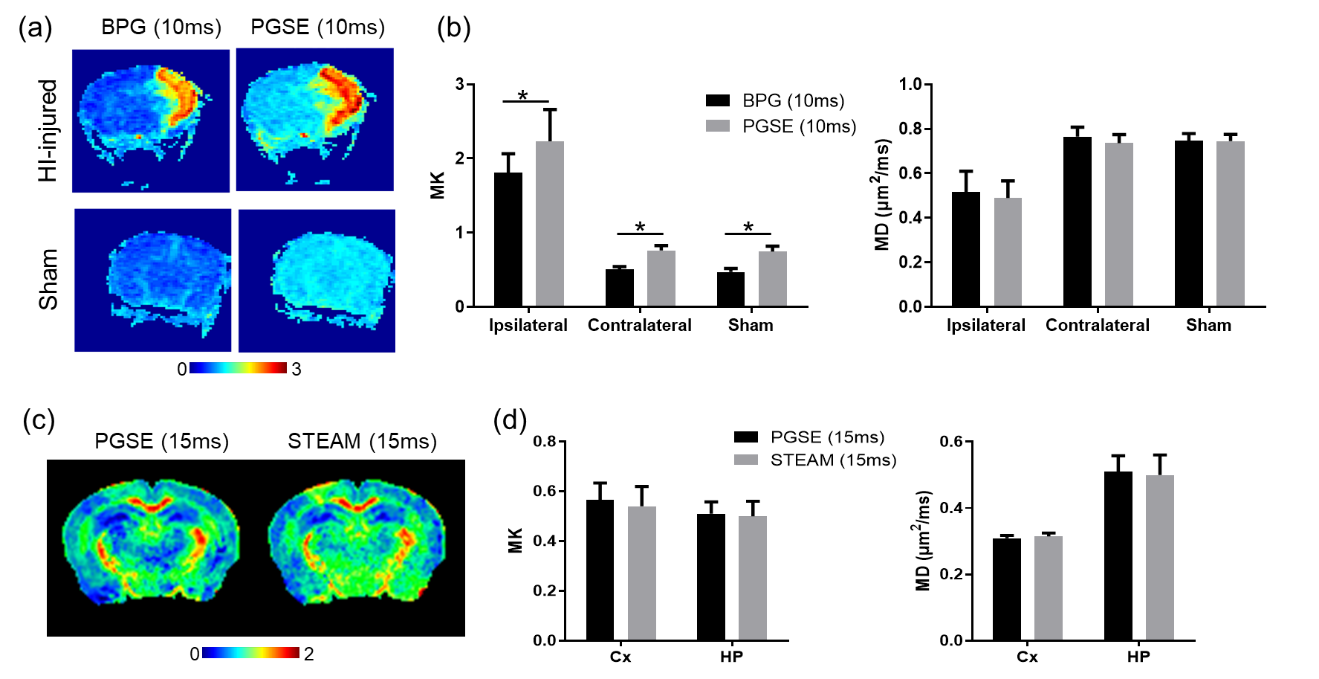

**Fig. S5:** Comparisons of different diffusion MRI encoding schemes. (**a**) BGP and PGSE based kurtosis maps at *t* of 10 ms in the HI-injured and sham mice. (**b**) Statistical comparison showed significantly lower (*p*<0.05, *n*=6) MK obtained from the BGP sequence than that from PGSE at 10 ms, while the MD were equivalent between the two sequences in both injured and sham mice. (**c**) PGSE and STEAM based kurtosis maps at *t* of 15 ms of a normal mouse brain. (**d**) Statistical comparison showed no difference (*p*>0.05, *n*=5) between the two sequences either in MK or MD between the two sequences.
